# Spatial Distribution of Cholesterol in Lipid Bilayers

**DOI:** 10.1101/636845

**Authors:** M. Aghaaminiha, S. Sharma

**Affiliations:** Ohio University

**Keywords:** Lipid bilayer, Cholesterol, Spatial distribution, Orientation, Molecular Simulation

## Abstract

1.

We have performed molecular simulations to study spatial distribution and orientation of cholesterol molecules within three symmetric lipid bilayer systems: two binary lipid mixtures, namely, DOPC/CHOL (1, 2-dioleoyl-sn-glycero-3-phosphocholine/cholesterol) and SM/CHOL (d20:1/20:0 sphingomyelin /Cholesterol), and a tertiary mixture of DOPC/SM/CHOL. We have studied the behavior of these bilayers at temperatures varying from 400 K to 210 K and cholesterol molar concentration from 0% to 60%.We observe that the spatial distribution of cholesterol is strongly correlated with the phase of the bilayer. In the *disordered* phase, cholesterol molecules are predominantly present near the center of the bilayer. In the *ordered* phase, cholesterol molecules are mainly present in the leaflets. At the *order* - *disorder* transition temperature, the fraction of cholesterol molecules is equal in the two leaflets and the center. In the leaflets, cholesterol molecules are oriented parallel to the bilayer normal, while near the center, cholesterol molecules are randomly oriented. In agreement with previous experimental studies, we find that increasing the cholesterol concentration favors *ordered* phase of the bilayers. The preference of cholesterol molecules to be present in the leaflets in the *ordered* phase is attributed to their favorable hydrophobic interactions with the lipid tails.

**Statement of Significance:** Cholesterol has an important role in governing the physical properties of lipid bilayers, including their structural integrity, phase behavior and permeability. The spatial distribution of cholesterol in lipid bilayers is not well-understood because of the challenges associated with performing experiments for such a measurement. We show, via molecular simulations, that the spatial distribution of cholesterol molecules is strongly correlated with the phase behavior of the lipid bilayers. In the *ordered* phase, cholesterol molecules are predominantly present in the leaflets, whereas in the *disordered* phase, cholesterol molecules are in the center region of the bilayer. These results are important for understanding the relationship between lipid bilayer composition and their biological function and response.

## 3. Introduction

Biological membranes act as boundaries to maintain internal organelles of cells safe from external environment (1–3). They form a semi-permeable interface that allows cells to interact with extracellular environment (2–5). In mammalian cells, biological membranes, also known as plasma membranes, are made up of proteins, lipids, sterols, and other molecules (1, 3). Proteins are thought to be responsible for the transport of molecules and for chemical signaling processes across plasma membranes (3, 6). The lateral mobility and function of membrane proteins is understood to be dictated by the structure of the membrane (6, 7). Cholesterols, the main sterol in plasma membranes, play an important role in the dynamics and structural properties of the membranes (1–3, 6, 7). Cholesterols affect bending modulus (8), and phase behavior of the membrane (9), and are important for the formation of rafts domains (10). In addition, cholesterols affect membrane fluidity, permeability (7, 11), lateral diffusivity (7), thickness, structural orientation (2, 7, 8, 10, 12), and lateral density (2, 5, 7) of lipids in the membrane. Since a lipid bilayer is the dominant constituent of plasma membranes, study of their behavior as a function of cholesterol concentration and temperature can provide useful insights about plasma membranes (5, 7, 9, 11, 13–15). However, within lipid bilayers, the spatial distribution of cholesterol has not yet been clearly established, mainly because of the difficulty associated with performing experiments for such a measurement (12).

Cholesterol solubility in a lipid bilayer is known to be governed by the size of head group of lipids as well as by lipids’ packing density (16, 17). Feigenson and Buboltz in 2001 reported that the maximum cholesterol concentration in the ternary mixture of DLPC/DPPC/CHOL is 66%, and above that, monohydrated crystals of cholesterol precipitate out from the bilayer (18). The head group of lipids acts as a shield to isolate cholesterols from the lipid-water interface (16, 18). Thus, in lipids with a larger head group, such as sphingomyelins, cholesterol has higher solubility. Furthermore, it is reported that cholesterol solubility in membranes composed of saturated lipids is significantly higher in comparison to in membranes with monounsaturated/unsaturated lipids (12, 16, 19).

Lipid bilayers are known to predominantly display two phases: *ordered* (liquid or gel) and *disordered*. These phases can be distinguished by considering translational order and chain configurational order of lipids in the bilayers (12, 16). The translational order refers to local structure of lipids and can be studied by calculating two dimensional radial distribution function (2D RDF) of lipid molecules in the bilayer plane. Chain configurational order is a measure of alignment of hydrocarbons tails of lipids in the bilayer. In the *ordered* phase, both translational order and chain configurational order are high, while in the disordered phase, both these orders are low. In the binary and ternary mixtures, cholesterol is well-known to favor the formation of the *ordered* phase ((9, 12, 13, 20), by enhancing the chain configurational order (12, 16, 21).

Molecular simulations have been useful in improving our understanding of biological processes, including the behavior of cell membranes (2, 22). While all-atom MD simulations have been employed to study dynamics and structural properties of lipid bilayers, these simulations are limited to time-scales of O(10^2^ ns) and length-scales of O(10 nm) (23, 24). Study of equilibrium phase behavior of lipid bilayers requires time-scales of O(1 μs) (7). Coarse-grained (CG) models reduce degrees of freedom of a system, and thus allow us to study longer length- and time-scales (23, 25). MARTINI force field is a popular CG force field for the study of lipid bilayers, surfactants, cholesterol, sugars and proteins (23, 26, 27). Along with the explicit solvent version (called Wet MARTINI), an implicit solvent version of the force field, known as Dry MARTINI has been recently developed (27). In the MARTINI force field, the coarse graining of molecules follows a four-to-one mapping, that is, each CG bead (or interaction site) represents four heavy atoms (23, 26). There are four main types of beads: polar, non-polar, apolar, and charged, along with several sub-types. In the Dry MARTINI force field, the non-bonded interactions matrix from wet MARTINI force field is re-parameterized to match the experimental values of hydration, vaporization, and partitioning free energies (27).

In this work, we have studied the spatial distribution and orientation of cholesterol molecules in DOPC/CHOL (1, 2-dioleoyl-sn-glycero-3-phosphocholine/cholesterol), SM/CHOL (d20:1/20:0 sphingomyelin /Cholesterol), and DOPC/SM/CHOL mixtures over a range of temperatures and cholesterol concentrations using CG MD simulations. Lipids and cholesterol molecules are modeled using the Dry MARTINI force field (see Figure 1). We have calculated structural and dynamical properties of the bilayers to identify different phases as a function of temperature and cholesterol concentration. We find that in the *ordered* phase, cholesterol is mainly present in bilayer leaflets, while in the *disordered* phase, cholesterol is predominantly present near the center of the bilayer. In the leaflets, cholesterol molecules are oriented parallel to the bilayer normal, whereas, cholesterol molecules near the center of the bilayer are randomly oriented with respect to the bilayer normal. Favorable hydrophobic interactions between the lipid tails and cholesterol molecules drives the preference of cholesterol to enter the leaflets.

**Figure 1:**
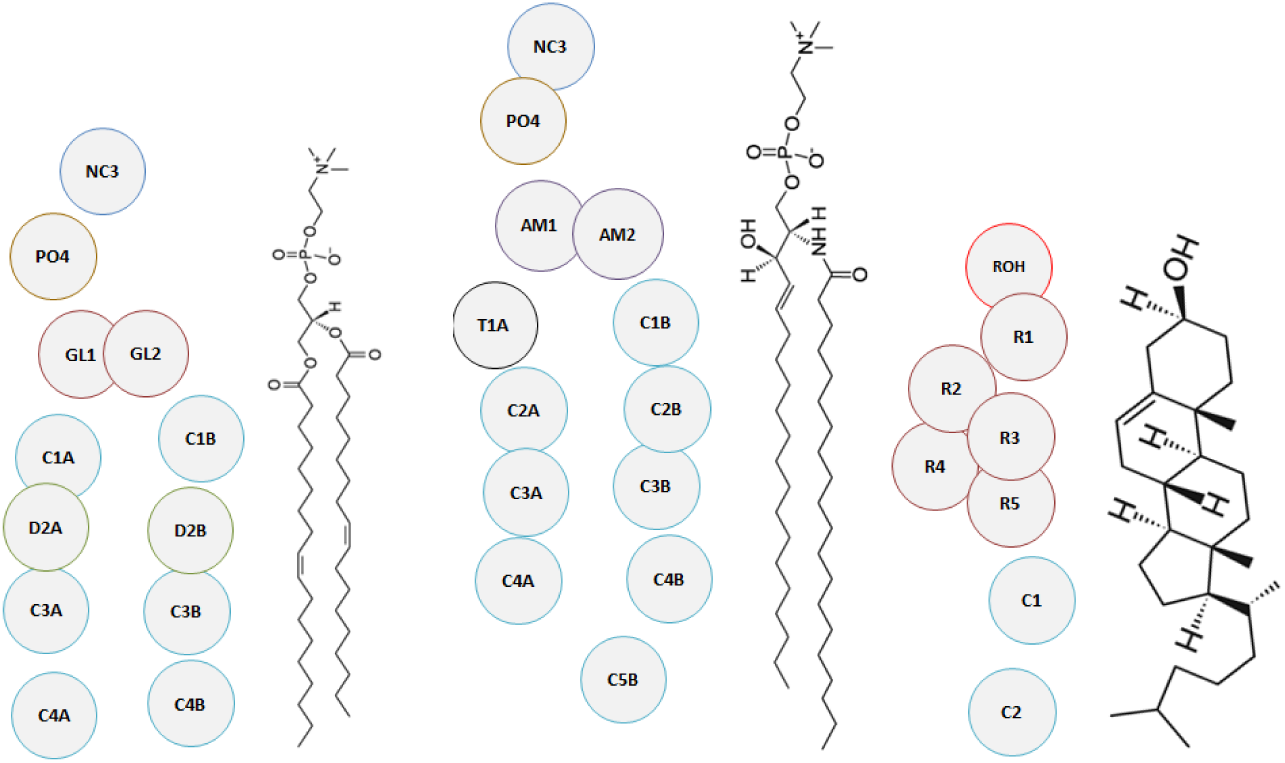
Dry Martini coarse grained beads. From left to right: DOPC, SM, and Cholesterol (28–30)

## 4. Simulation System and Methods

We studied three different types of lipid bilayers: DOPC/CHOL, SM/CHOL, and 1:1 DOPC/SM/CHOL. DOPC represents a lipid with low melting temperature. SM represents a lipid with relatively high value of melting temperature. All lipids and cholesterol molecules were modeled using Dry MARTINI version v2.0 force field. Periodic boundary conditions were applied in all three directions of the simulation box. All simulations were performed using the GROMACS/5.1.2 (31) molecular simulation package. The initial configuration of the three bilayer systems was constructed using the *insane.py* script developed by Marrink et al (28). The total number of molecules (lipids and cholesterols) were similar in all the simulations (∼512, kindly see Table S1, Supporting Information for the exact numbers). The concentration of cholesterol in the lipid bilayers was varied from 0% to 60% mole-ratio.

### 4.1. Simulation Protocol

After generating the initial configuration of the lipid bilayer, the energy of the configuration was minimized using the steepest descent algorithm. Next, to equilibrate the bilayer structure, a 50 ns NVT ensemble simulation followed by a 50 ns NPT ensemble simulation was performed at a constant temperature of 400 K. The time-step for the simulations was 10 fs. After the simulation at 400 K, the system was cooled to 390 K and a 10 ns equilibration followed by a 40 ns of production run was performed. This way, the entire temperature range from 400 K to 210 K was studied. For the three bilayer systems at temperatures < 300 K, we performed 500 ns of production run to ensure invariability of results. For the three bilayers, we also studied systems with four times the number of molecules for few temperatures to ensure there are no finite size effects in the results. Figure 2 shows the initial configuration of DOPC/SM/CHOL system before performing MD production run at 400 K.

**Figure 2:**
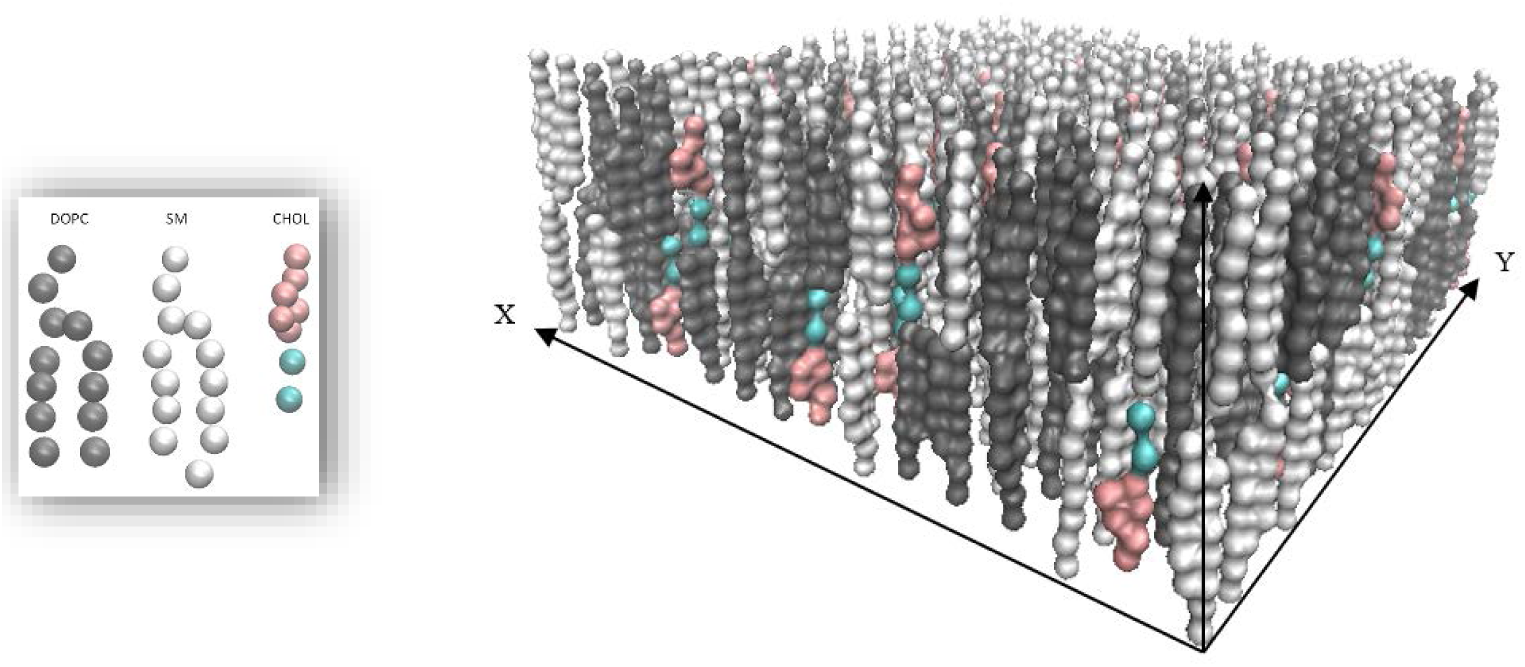
Mixture of DOPC/SM/CHOL initial configuration (at molar cholesterol concentration of 10%)

### 4.2. Simulation parameters

Spherical cut-off for Lennard-Jones interactions was taken as 1.2 nm. The non-bonded forces were smoothly switched to zero beyond the spherical cut-off. Velocity-rescaling thermostat developed by Bussi et al (32) was used as the temperature coupling method. The time constant for the thermostat was taken to be 1 ps. For the NPT runs, Parrinello-Rahman barostat was employed with the time constant of 4 ps. In the NPT runs, the membrane was made tensionless by setting the reference pressure to 0 bar in *xy*-plane. Also, the isothermal compressibility was set to 3 × 10^−4^ bar^−1^ in the *xy*-plane and 0 bar^−1^ in *z* direction (27). To remove net motion of the bilayer, the center of mass motion was subtracted out.

### 4.3. Bilayer properties analysis

Area per lipid was calculated as the ensemble-averaged area of the plane in the simulation box in which the bilayer resides divided by the number of lipids in one leaflet. Bilayer thickness was calculated as the ensemble-averaged distance between the PO_4_ beads (see Figure 1) of lipids in each leaflet. The average tail angle with bilayer normal was calculated as the angle between vector-a (vector connecting the first bead to the last bead of each tail, see Figure 1) and vector-b (z axis). To monitor ordering of lipid chains we calculated the ensemble-averaged order parameter as follows:

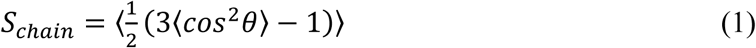

Where θ is the angle between each bond vector of the CG lipid beads and the bilayer normal (*z*-axis). High value of the *S*_*chain*_ (∼ 1) indicates that the bonds are aligned with the bilayer normal, representing the chain order characteristic of the *ordered* phase (10, 16) while its low value (∼ 0) indicates that the bonds are randomly oriented with respect to the bilayer normal, a signature of the *disordered* phase (10, 16). To quantify translational ordering of lipids, we calculated two dimensional (2D) radial distribution function (RDF). Long-ranged and more ordered correlations belong to the *ordered* phase while short-ranged correlations that decay rapidly represent the *disordered* phase. We calculated the 2D RDF based on the C1B beads (see Figure 1) of lipids, similar to (1) since it is located in the middle of both DOPC and SM and can be taken as the center of mass of the lipids. Phase behavior of lipid bilayers has been extensively studied before, and different phases have been classified as *gel, liquid-ordered* and *liquid-disordered* etc. Through heuristic rules to identify these phases, previous simulation studies have developed phase diagrams of lipid bilayers. In this work, our focus is to study spatial distribution of cholesterol in lipid bilayers as a function of the structure of lipid bilayers. Therefore, we have classified the behavior of lipid bilayers more broadly in terms of *ordered* and *disordered* phases only.

In addition to the structural properties mentioned above, we calculated the lateral diffusion coefficients of lipids in the bilayers. The angle of cholesterol molecules with the bilayer normal (*z*-axis) was also calculated as the ensemble-averaged of the angle between vector-a (vector connecting ROH bead to C2 bead, see Figure 1) and vector-b (z axis).

## 5. Results and Discussion

To determine spatial distribution of cholesterol molecules within the lipid bilayer and find its dependence on the phase of the bilayer, we studied three lipid/cholesterol mixtures, namely, DOPC/CHOL, SM/CHOL, and DOPC/SM/CHOL. For each system, we studied bilayer properties by varying the temperature from 400 K to 210 K in decrements of 10 K for seven different cholesterol concentrations ranging from 0% to 60%.

### 5.1. Behavior of lipid bilayers as a function of temperature in the absence of cholesterol

As a first step, we calculated structural properties of lipid bilayers of pure DOPC, pure SM and 1:1 mixture of DOPC/SM in the absence of any cholesterol. Figure 3 shows area per lipid, average bilayer thickness, average order parameter (*S*_*chain*_), and 2D RDF. Other structural and dynamical properties of bilayers such as average tail angle, total energy, and diffusivity are presented in Figure S1 (Supporting Information). From structural properties (Figure 3 A to C) as well as from 2D RDFs (Figure 3 D to F), it is observed that pure SM and 1:1 mixture of DOPC/SM have significant ordering of lipids at low temperatures, and as the temperature increases, they undergo an *order* - *disorder* transition. In contrast, pure DOPC bilayer does not show significant ordering even at the lowest temperatures sampled. This behavior of DOPC lipids matches with experimental observations as well (14). To estimate the transition temperature, *T*_*m*_ of pure SM and 1:1 mixture of DOPC/SM systems, we fitted a polynomial function to the area per lipid graph and determined the location of the inflexion point by setting the second derivative to zero. Figure S2 (Supporting Information) shows plot of the derivative of the fitted polynomial. The *T*_*m*_ of pure SM is estimated to be 315 K, which agrees the experimentally reported value of 313 K (13). The *T*_*m*_ of 1:1 mixture of DOPC/SM system is estimated to be 250 K. Interestingly at the *T*_*m*_ for both SM and DOPC/SM systems, *S*_*chain*_ ∼ 0.5. Furthermore, the 2D RDFs presented in Figure 3 (E and F) do show loss of long-range structure as the temperature increases beyond the *T*_*m*_. For DOPC, the *S*_*chain*_ never goes above 0.3, and the 2D RDFs do not show any long-range structure, supporting the conclusion that the DOPC lipid bilayer does not have ordered structure at any of the sampled temperatures.

**Figure 3:**
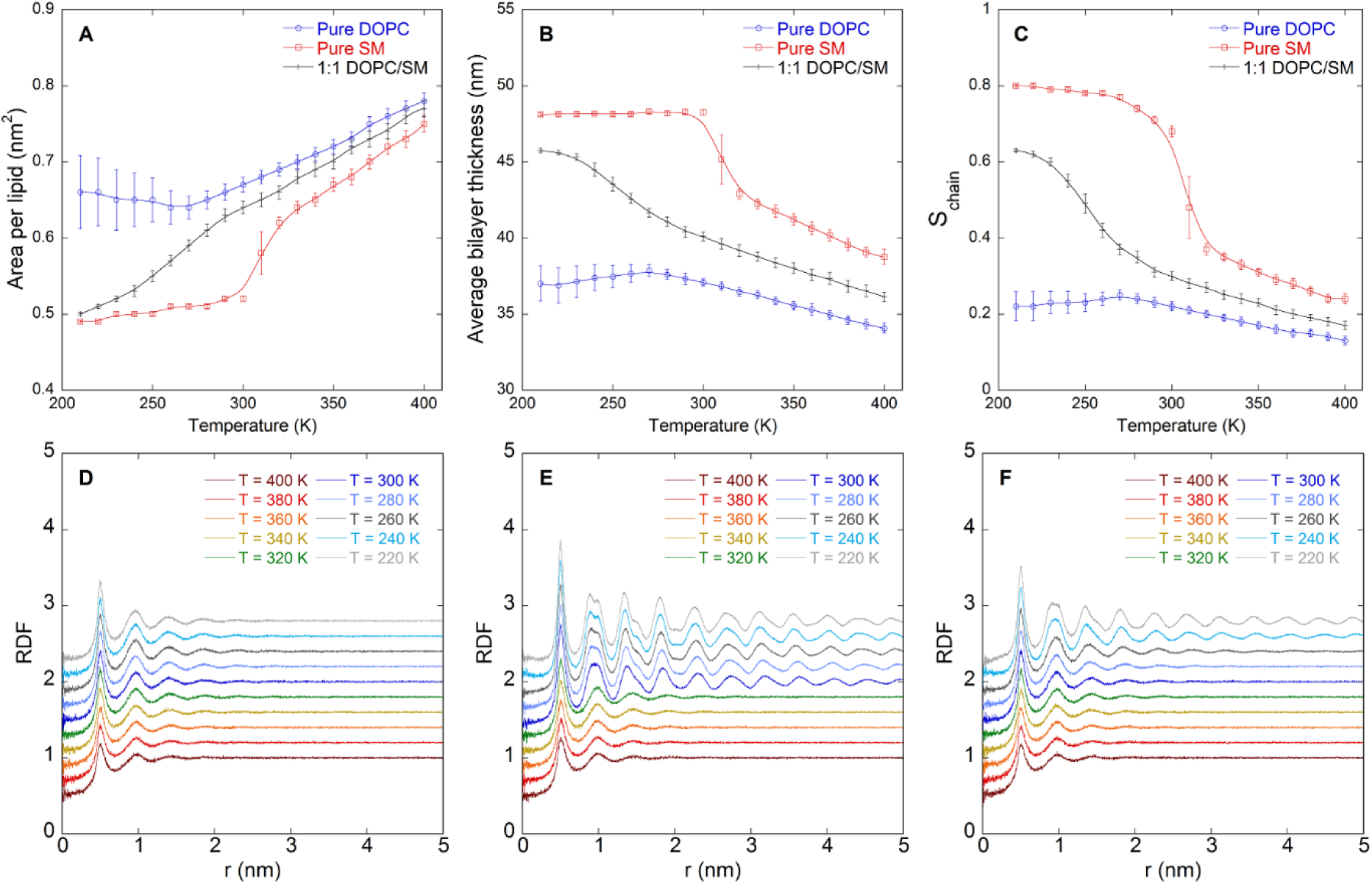
Bilayer structural properties as a function of temperature: Area per lipid (A), average bilayer thickness (B), average order parameter (*S*_*chain*_) (C), 2D RDF of pure DOPC (D), pure SM (E), and 1:1 DOPC/SM (F). Successive 2D RDFs have been shifted vertically by 0.2 units for the sake of clarity

### 5.2. Spatial distribution of cholesterol in lipid bilayers

We performed molecular simulations of the three lipid bilayer systems with molar concentration of cholesterol varying from 10% to 60% using the simulation protocol discussed above. Figure 4 shows fraction of cholesterol molecules in the upper leaflet, lower leaflet and center of the lipid bilayers when the overall molar concentration of cholesterol is 10%. The details of how upper leaflet, lower leaflet and center regions are defined are provided in Figure S3 (Supporting Information) and the accompanying text. From Figure 4, it is seen that in the case of DOPC, a larger fraction of cholesterol molecules are in the center region for all temperatures. However, for pure SM and 1:1 DOPC/SM mixture, there is higher fraction of cholesterol molecules in the upper and lower leaflets at lower temperatures. As temperature increases, the fraction of cholesterol molecules in the center region becomes higher. The crossover occurs close to the *T*_*m*_ for these two lipid bilayer systems. Figure S4 (Supporting Information) shows the fraction of cholesterol molecules in the three lipid bilayer systems for molar cholesterol concentrations ranging from 20% to 60% as a function of temperature. It is observed that for all concentrations and temperatures, the cholesterol fraction is higher in the center region in the DOPC lipid bilayer. On the other hand, the pure SM and 1:1 DOPC/SM mixture show a crossover in the fraction of cholesterol molecules between the regions. The temperature at which the fraction of cholesterol molecules in the leaflet and the center regions becomes equal is denoted as *T*_*cross*_.

**Figure 4:**
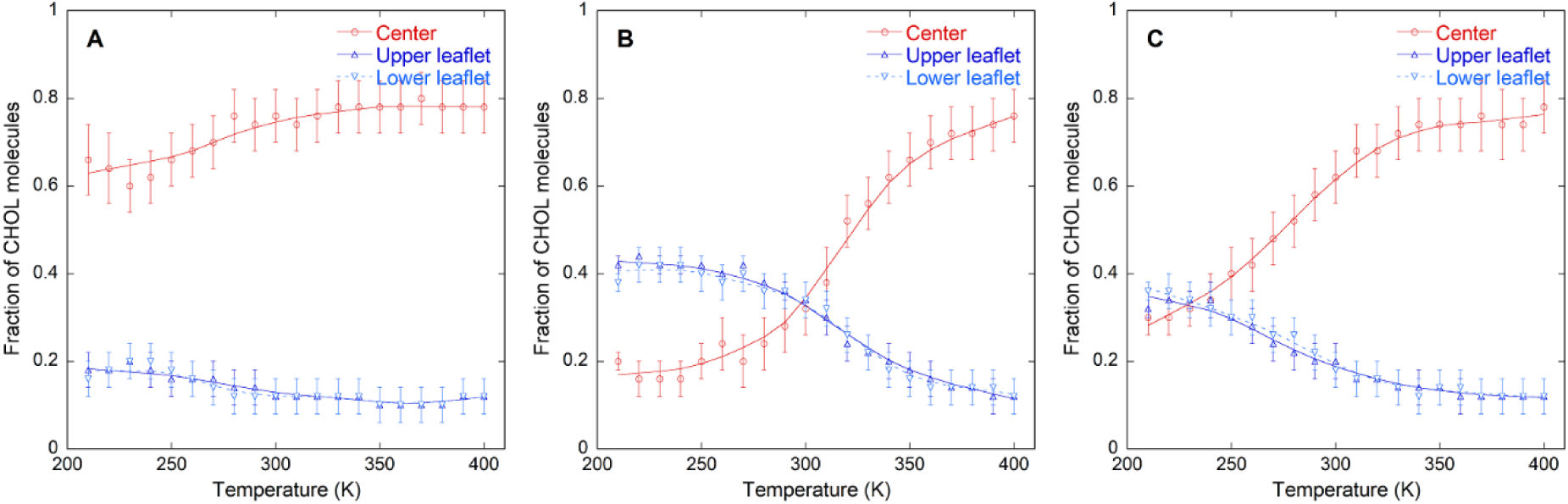
Fraction of cholesterol molecules at different regions of the bilayer (at molar cholesterol concentration of 10%). Red, dark blue, and light blue lines represent cholesterol molecules located at the center, upper leaflet, and lower leaflet, respectively. DOPC/CHOL (A), SM/CHOL (B), and DOPC/SM/CHOL (C)

Figure 5 shows the *S*_*chain*_ parameter for the three lipid bilayer systems for different cholesterol concentrations as a function of temperature. The *order* - *disorder* transition, *T*_*m*_, for each system is denoted as the temperature at which *S*_*chain*_ = 0.5. This definition of *T*_*m*_ is employed because it is always found to be within a few Kelvins of the *order-disorder* transition temperature determined by locating the point of inflexion in structural properties of lipid bilayers (see Table S3, Supporting Information). Clearly from Figure 5A, the DOPC lipid bilayer is always in the *disordered* phase. For both SM and 1:1 DOPC/SM mixture, the *T*_*m*_ shifts towards higher values as the concentration of cholesterol increases. The 2D RDFs of these systems are shown in Figures S6 (DOPC/CHOL), S7 (SM/CHOL) and S8 (DOPC/SM/CHOL) in Supporting Information. The shift in *T*_*m*_ to higher values with increasing cholesterol concentration have also been reported for the system of DPPC/CHOL (7).

**Figure 5:**
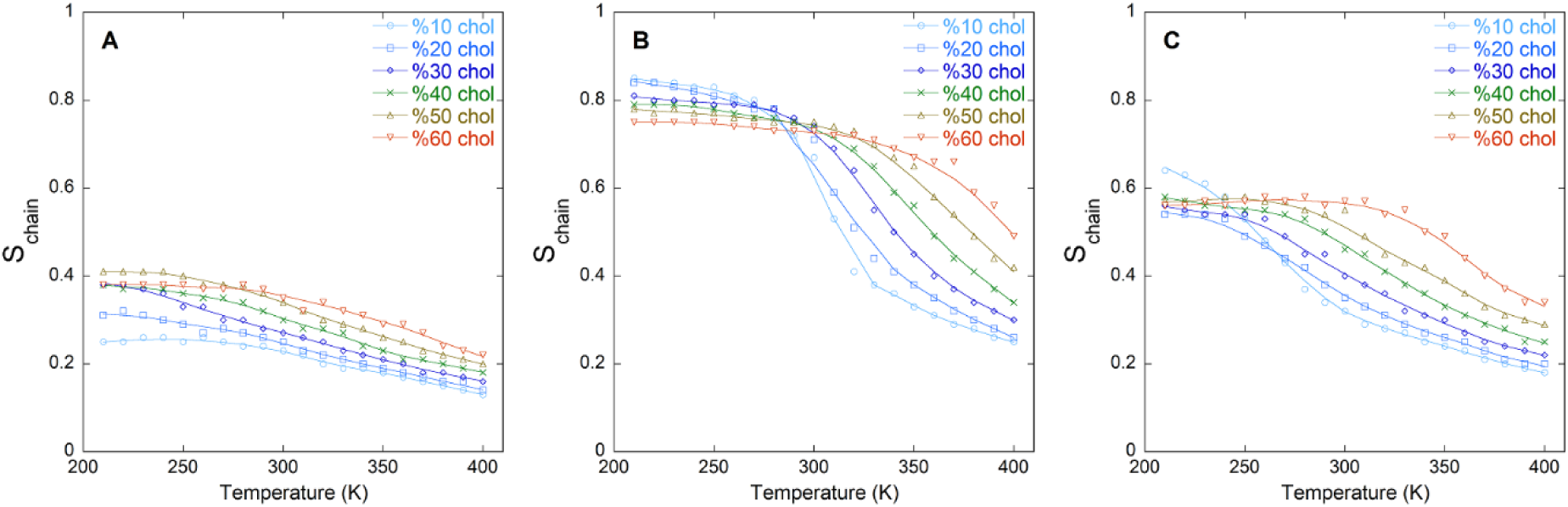
Average order parameter of chains (*S*_*chain*_) in the three systems: DOPC/CHOL (A), SM/CHOL (B), and DOPC/SM/CHOL (C)

Figure 6A shows a plot of *T*_*m*_ and *T*_*cross*_ for the pure SM and DOPC/SM systems as a function of cholesterol concentration. Figure 6B shows a comparison of *T*_*m*_ and *T*_*cross*_. The *T*_*m*_ and *T*_*cross*_ increase with cholesterol concentration, as expected. Also, the *T*_*m*_ and *T*_*cross*_ lie close to the *y* = *x* line in Figure 6B indicating that the spatial distribution of cholesterol is strongly correlated with the ordering in lipid bilayers.

**Figure 6:**
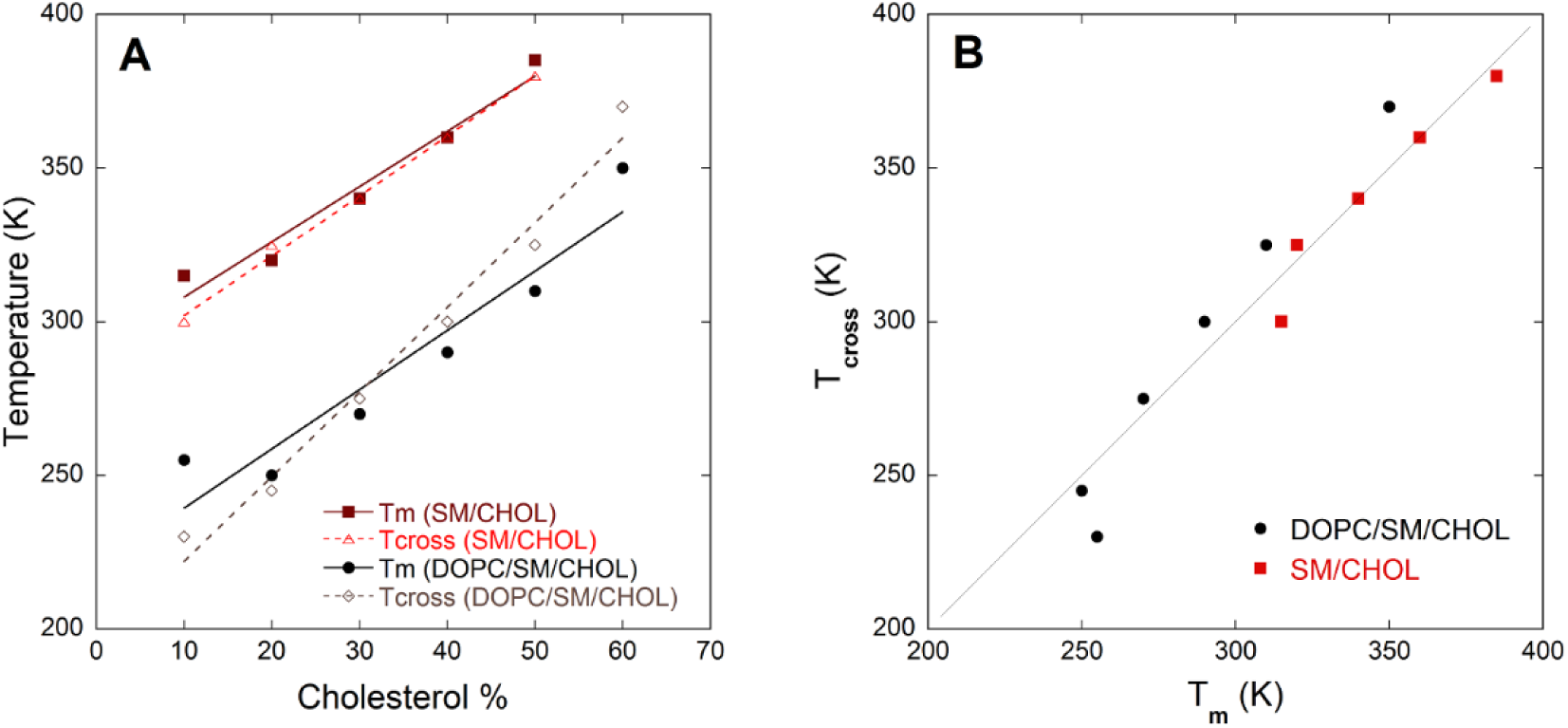
Spatial distribution of cholesterol vs phase represented with average order parameter (S_chain_)

According to Figure 4 and Figure S4 (Supporting Information), the DOPC/CHOL lipid bilayer system does not show any *T*_*cross*_ and cholesterol always prefers to be in the center region. This matches with the observation that the DOPC/CHOL lipid bilayer is always in the *disordered* phase. The result that the DOPC system behaves differently from the other two systems is in agreement with the previous studies that have investigated the affinity of cholesterol in different types of lipids. In these studies, it is concluded that unsaturated fatty acids including lipids such as DOPC, have less affinity for cholesterol due to high disorder in these bilayers, however saturated lipids such as SM, have more favorable interactions with cholesterol (12, 13).

Figure 7 shows density profiles of cholesterol molecules at different temperatures and for 10% molar cholesterol concentration for the three lipid bilayer systems studied. These profiles support our conclusion that cholesterol molecules prefer to be located at the center of the bilayer at high temperatures when the bilayer is in the *disordered* phase. At low temperatures for pure SM and 1:1 DOPC/SM mixture bilayers, cholesterol molecules prefer to be in the leaflets when the bilayer is in the *ordered* phase. Cholesterol is a hydrophobic molecule. In the *ordered* phase, it is able to form significant hydrophobic interactions with lipid tails. Hence, the entropic loss of entering the leaflets is compensated by the energetic gain of hydrophobic interactions. Figures S9 to S13 (Supplemental Information) display density profiles of the three systems at higher cholesterol concentrations (20% to 60%).

**Figure 7:**
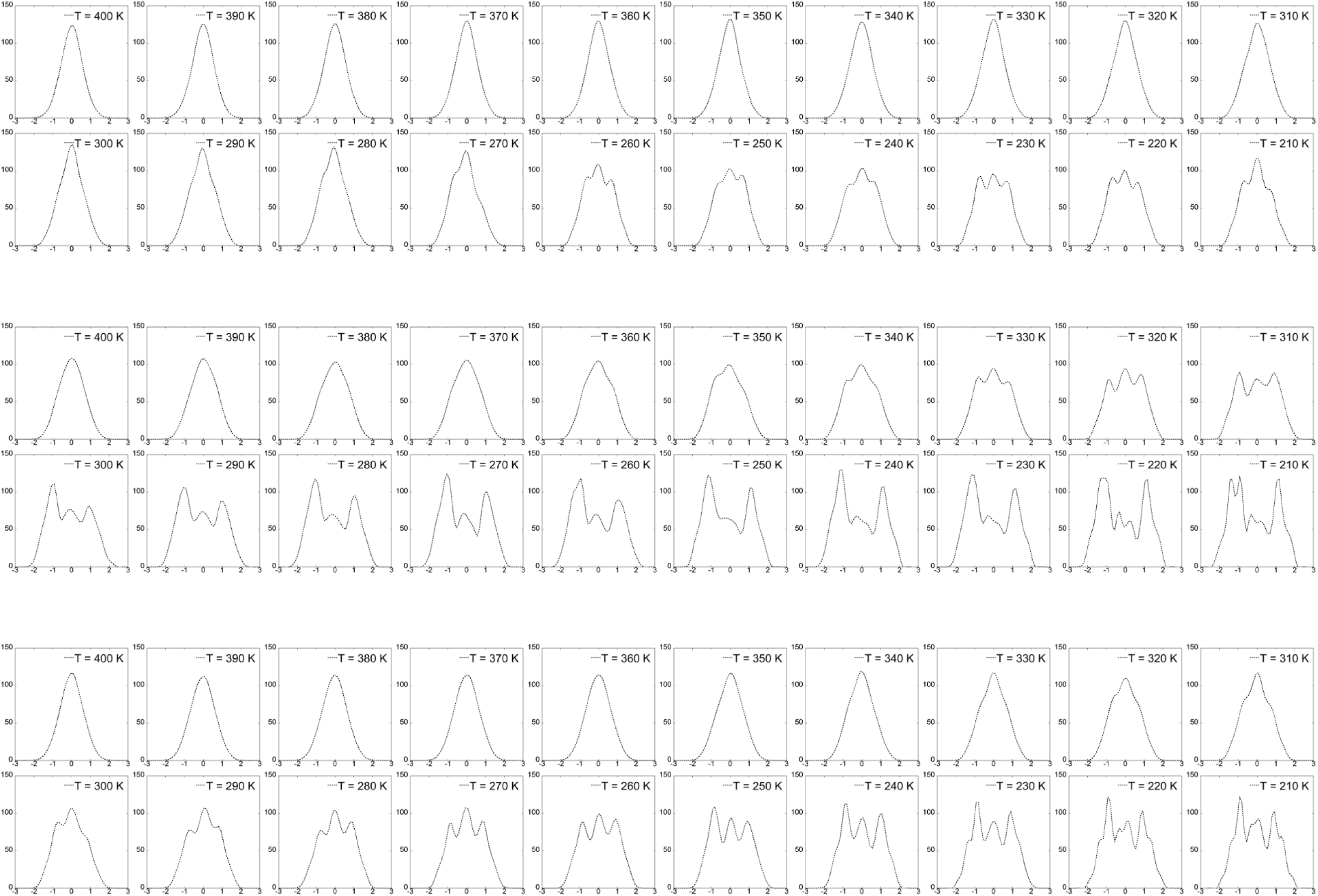
Cholesterol density profile at molar cholesterol concentration of 10%. (Top) DOPC/CHOL, (middle) SM/CHOL, (bottom) DOPC/SM/CHOL In all plots, *y* axis is the density of cholesterol in *g/L* from 0 to 150; and *x* axis is average relative position from center of the bilayer in *nm* from −3 to +3.

Figure 8 shows snapshots of cholesterol molecules in the three lipid bilayers at 10% molar cholesterol concentration and at two temperatures 400 K and 210 K. The snapshots show that SM/CHOL and DOPC/SM/CHOL lipid bilayers have ordered cholesterol molecules at 210 K, while the DOPC/CHOL bilayer does not show any significant ordering.

**Figure 8:**
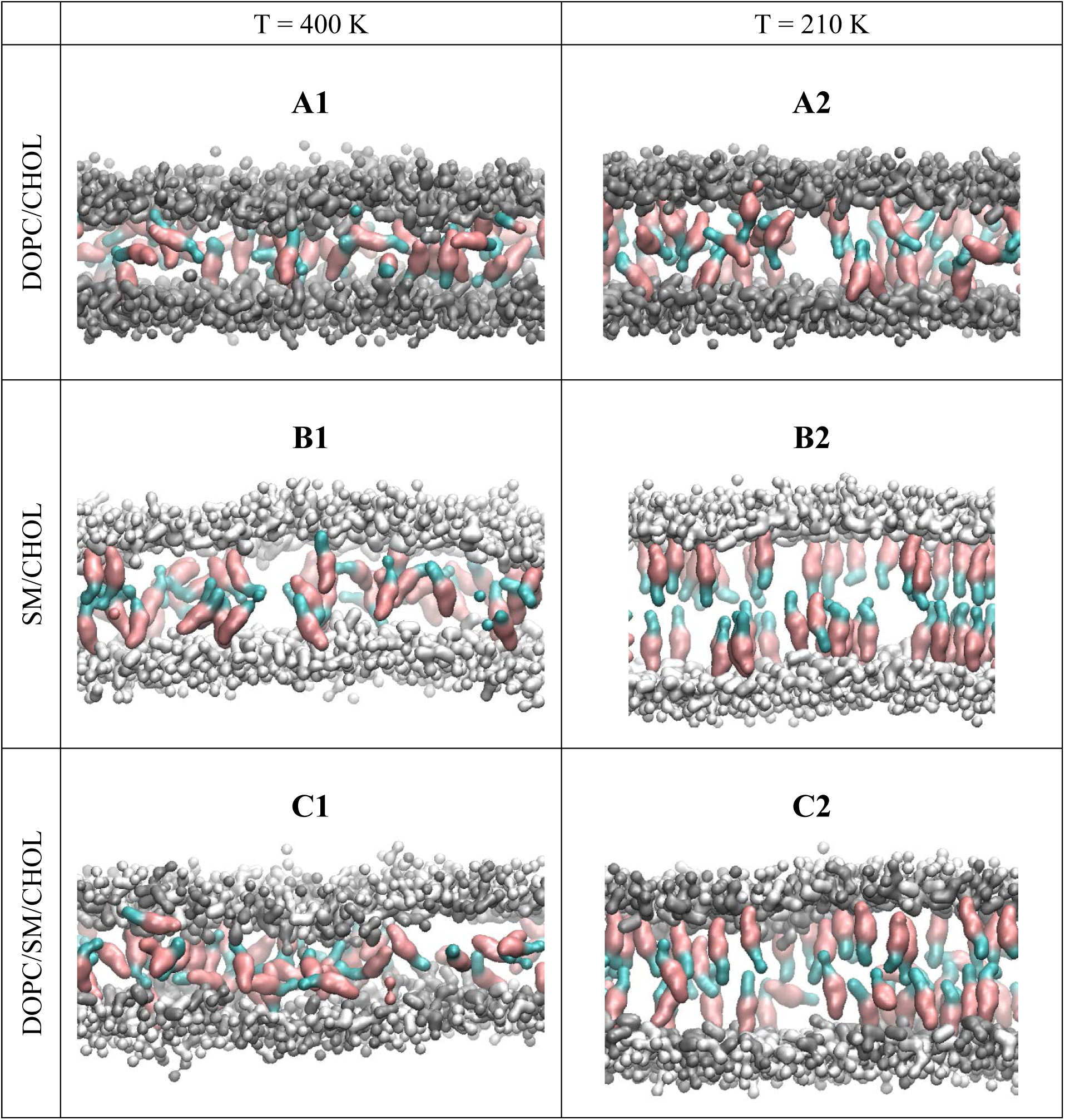
Simulations snapshots at 400 K and 210 K for the three systems (at molar cholesterol concentration of 10%). The grey and white beads represent SM and DOPC head groups respectively, the maroon and cyan colors represent hydrocarbon rings and hydrocarbon tail of cholesterol, respectively. At T = 400 K (Figure A1, B1, and C1), the thickness of the bilayers is small, cholesterol molecules are mainly at the center, and the orientation of cholesterol molecules is random. At T = 210 K and for SM/CHOL and DOPC/SM/CHOL systems, (Figures B2 and C2), cholesterol molecules are oriented parallel to the bilayer normal. Tails of lipids are omitted in these snapshots for the sake of clarity.

### 5.3. Orientation of cholesterol molecules

Figure 9 shows average angle that cholesterol molecules in the center region and leaflets make with the bilayer normal (*z* axis) when the molar cholesterol concentration is 10%. In SM/CHOL and DOPC/SM/CHOL bilayers, the cholesterol molecules are more aligned with the bilayer normal at low temperatures. The average angle in the leaflets is smaller than at the center, implying that cholesterol molecules align parallel to the bilayer normal in the leaflets. The average angle of cholesterol molecules in the three bilayers for higher cholesterol concentrations (20% to 60%) are shown in Figure S14 (Supporting Information). Figure 9 also shows that cholesterol molecules in the DOPC/CHOL system are less oriented with the bilayer normal as compared to the other two systems.

**Figure 9:**
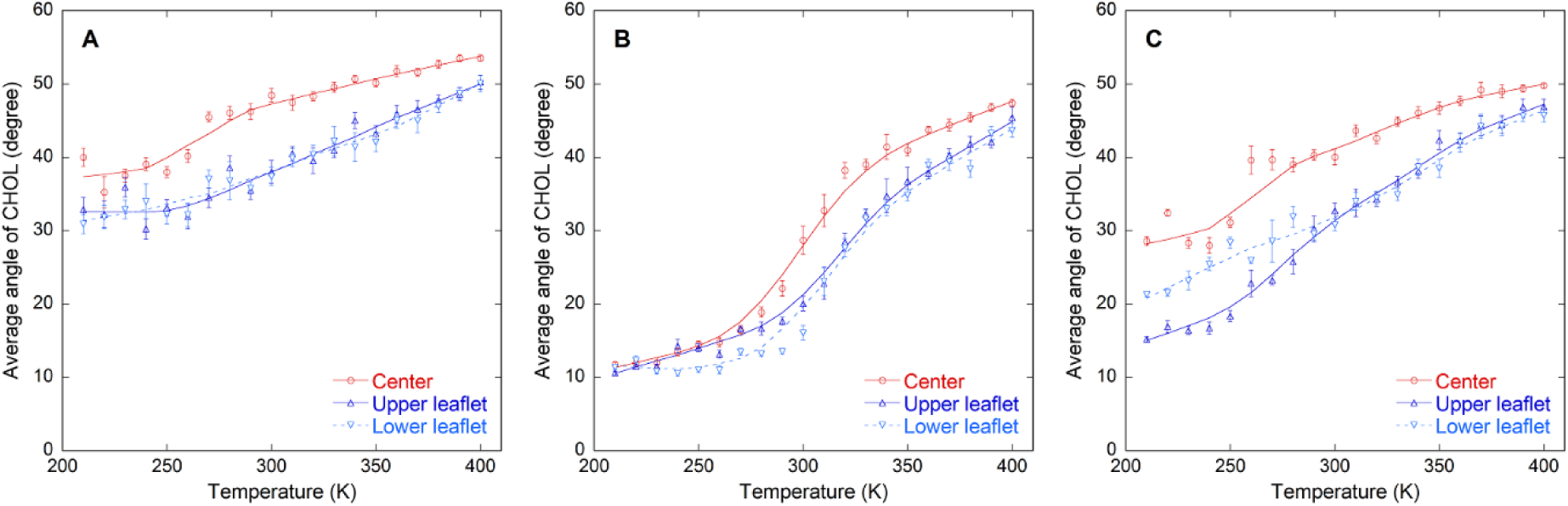
Orientation of cholesterol molecules in different regions of the bilayer (at molar cholesterol concentration of 10%). Red, dark blue, and light blue lines represent cholesterol molecules located at the center, upper leaflet, and lower leaflet, respectively. DOPC/CHOL (A), SM/CHOL (B), and DOPC/SM/CHOL (C)

## 6. Conclusions

We have studied spatial distribution of cholesterol molecules as a function of cholesterol concentration and temperature in three lipid bilayer systems: DOPC/CHOL, SM/CHOL and DOPC/SM/CHOL using coarse-grained molecular simulations. We find that the spatial distribution of cholesterol molecules is closely related to the *ordered* and *disordered* phase of the bilayers. In the *ordered* phase, cholesterol molecules prefer to be in the leaflets and align themselves parallel to the surface normal. In this configuration, the cholesterol molecules maximize their hydrophobic interactions with the lipid tails. In the *disordered* phase, cholesterol molecules are preferentially in the center region of the bilayer. DOPC/CHOL bilayer does not form an *ordered* phase in the sampled temperature range. Therefore, for this system, the cholesterol molecules are always found predominantly in the center region. The temperature of the cross-over, *T*_*cross*_ defined as the temperature at which the fraction of cholesterol molecules in the leaflets is equal to the fraction in the center region is found to be quite close to the *order* - *disorder* transition temperature, *T*_*m*_ for all the bilayer systems studied.

## 7. Authors Contribution

MA and SS designed research. MA performed research. MA and SS performed analysis of simulation data. MA and SS wrote the manuscript.

## 8. Acknowledgements

The authors would like to acknowledge Ohio University’s start-up funds for covering the computational costs of performing simulations at the Ohio Supercomputer Center. SS acknowledges additional computational resources from National Science Foundation XSEDE program (request number DMR190005).

